# Antibody responses to immunization require sensory neurons

**DOI:** 10.1101/860395

**Authors:** Aisling Tynan, Téa Tsaava, Manojkumar Gunasekaran, Isabel Snee, Tak W. Mak, Peder Olofsson, Ulf Andersson, Sangeeta S. Chavan, Kevin J. Tracey

## Abstract

Mammals store memories in the nervous and immune systems. Sensory neurons have been implicated in enhancing neurological memory, but whether neurons participate during immunity to novel antigens is unknown. Here, mice rendered deficient in transient receptor potential vanilloid 1 (TRPV1)-expressing sensory neurons, termed “nociceptors,” fail to develop competent antibody responses to KLH and hapten-NP. Moreover, selective optogenetic stimulation of TRPV1 neurons during immunization significantly enhanced antibody responses to antigens. Thus, TRPV1 nociceptors mediate antibody responses to novel antigen, and stimulating TRPV1 nociceptors enhances antibody responses during immunization. This is the first genetic and selective functional evidence that nociceptors are required during immunization to produce antigen-specific antibodies.

**Summary:** The first genetic and selective functional evidence showing that TRPV1-expressing nociceptors are required for competent antibody responses to novel antigen, and stimulating TRPV1 nociceptors enhances antibody responses to novel antigen.

## Introduction

Natural selection conferred mammals with the exquisite capacity to retain information in two organ systems, nervous and immune. These principle sensory systems recognize, integrate and respond to various internal and external stimuli. Both systems can recall previous exposure to stimulli, and develop memories of these challenges, leading to efficient future “adaptive” responses. The nervous system responds to a broad spectrum of external and internal stimuli, and controls homeostasis through central and peripheral neuronal networks. The immune system surveys the internal environment, and defends against infection and injury through its tissue resident and circulating immune cells. Although the immune and nervous systems clearly perform unique functions, they communicate and regulate each other to maintain homeostasis. Several studies have identified bidirectional crosstalk between nervous system and immune system in healthy state and pathologies in numerous conditions, including chronic inflammatory disorders, acute inflammatory conditions, autism, multiple sclerosis, and cancer (Louveau et al., 2015; Baral et al., 2018; Veiga-Fernandes and Mucida, 2016; Godinho-Silva et al., 2019). Recent studies have also revealed that specific anatomical locations of neuro-immune interactions may serve as important immunoregulatory hubs (Huang et al., 2019; Godinho-Silva et al., 2019; Veiga-Fernandes and Pachnis, 2017).

The nervous system perceives the environment by receiving input from sensory neurons, termed “nociceptors.” These specialized neurons densely innervate peripheral tissues, including the skin, joints, respiratory, and gastrointestinal tract, and detect various noxious stimuli such as chemicals, temperature and pathogens (Baral et al., 2018, 2019; Chiu et al., 2012b). Nociceptor sensory neurons share many molecular recognition receptor pathways as immune cells, which enables them to detect pathogens and other inflammatory signals, and directly communicate with the immune system in the event of danger (Lai et al., 2017; Chiu et al., 2013; Hanes et al., 2016; Gunasekaran et al., 2018). Bacteria, danger associated molecules such as ATP, inflammatory cytokines including IL1 and TNF can directly activate nociceptors leading to downstream signaling events (Chiu et al., 2013; Steinberg et al., 2016; Zanos et al., 2018; Cockayne et al., 2000; Mariathasan et al., 2006; Binshtok et al., 2008; Zhang et al., 2011; Samad et al., 2001). Upon activation, nociceptors release neuropeptides and neurotransmitters that directly attract and regulate innate immune cells (mast cells, dendritic cells, innate lymphoid type 2 cells) and adaptive immune cells (T lymphocytes) during the initial stages of infection or injury (Talbot et al., 2015; Pinho-Ribeiro et al., 2018; Wilson et al., 2013; Cyphert et al., 2009; Mikami et al., 2011; Rochlitzer et al., 2011; Ding et al., 2008). This bidirectional communication between the nociceptors and immune system cells have been studied in bacterial infection (Chiu et al., 2013), necrotizing fasciitis (Pinho-Ribeiro et al., 2018), atopic dermatitis (Riol-Blanco et al., 2014; Caterina et al., 2000), psoriasis (Riol-Blanco et al., 2014), and allergic airway inflammation (Talbot et al., 2015).

Activation of nociceptive sensory receptors also mediates the sensory experience of pain (Chiu et al., 2012b). Exposure to mild heat-induced pain significantly enhanced memory to recall pictures of everyday objects, and enhanced fMRI activity in the brain insula correlated with memory for pain-associated objects (Wimmer and Buchel, 2015; Liberzon and Martis, 2006). Indeed, hyperalgesia, pain sensitiviy, frequently results from injuries or inflammation of peripheral tissues (Sandkühler, 2009). While these studies implicate nociception in establishing memory in the nervous system, to our knowledge whether nociception participates in establishing immunological memory is unknown.

In 1989, Charles Janeway posited that mechanisms of immunological memory, also termed “adaptive immunity,” required genome encoded pattern recognition receptors sensitive to highly conserved molecules derived from microbial pathogens (Janeway, 1989). His theory, since proven, predicted that the evolutionarily ancient innate immune system is required for the competent storage of memory in cells of the adaptive immune system. Recently, we suggested a new revolution is at hand, one that requires understanding how nervous system mechanisms mediate adaptive immunity (Tracey, 2015). Thus, neurons that sense and respond to pathogenic challenges may be essential to competent storage of memory in cells of the adaptive immune system.

A prominent subset of nociceptor neurons express transient receptor potential vanilloid 1 (TRPV1), first identified as a receptor for capsaicin, the vanilloid isolated from chili peppers that is experienced as heat (Chiu et al., 2012b; Baral et al., 2019). TRPV1 activation of peripheral sensory neurons activates depolarizing sensory signals that propagate to the central nervous system which produces pain during infection and inflammation. TRPV1 activation also stimulates axon reflexes, in which nociceptors act with local efferent signaling arcs to release neurotransmitters, including substance P and calcitonin gene-related peptide (CGRP) in nearby tissues (Pinho-Ribeiro et al., 2018; Szallasi and Blumberg, 1999). TRPV1 sensory neurons play a major role as a polymodal nociceptor integrating physical and chemical stimuli, including low pH, lipid metabolites, exogenous and endogenous vanilloid compounds, and noxious heat (Szallasi and Blumberg, 1999; Helyes et al., 2007). Dysregulated TRPV1 function, through mechanisms mediated by local efferent secretory function, has been implicated in arthritis, colitis, Type 1 diabetes and lung inflammation (Helyes et al., 2007; Razavi et al., 2006; Szabó et al., 2005; Kimball et al., 2004).

Because nociceptors evolved to detect threat agents, we reasoned that TRPV1+ nociceptor neurons may regulate antibody responses to immunization. Here, silencing TRPV1-expressing nociceptor neurons in mice exposed to a novel antigen significantly reduced specific antibody responses, and selective optogenetic stimulation of TRPV1-expressing nociceptor neurons significantly enhanced specific antibody responses.

## Results

### TRPV1+ sensory neurons control onset of antigen-specific antibody responses

To understand the role of TRPV1-expressing nociceptors in modulating antigen-specific antibody responses, we pursued two different complementary strategies. First, we utilized a genetic approach to selectively ablate TRPV1-expressing nociceptors by crossing TRPV1-Cre mice that expressed Cre recombinase under the control of *TRPV1* locus, with floxed diphtheria toxin A (DTA) mice (Chiu et al., 2013). TRPV1-driven expression of DTA induces cell death by catalyzing the inactivation of elongation factor 2 and inhibiting protein synthesis (Collier, 2001; Maxwell et al., 1986; Palmiter et al., 1987). We have previously established that TRPV1-Cre/DTA mice do not express TRPV1+ neurons (Zanos et al., 2018).

Naïve TRPV1-ablated and wild-type control mice were immunized with keyhole limpet hemocyanin (KLH), a naturally occurring respiratory protein of giant keyhole limpets living in a sea, which is not ordinalrily encountered by mammalian immune system (Harris and Markl, 1999; Kantele et al., 2011). KLH-specific antibody responses were monitored in serum at day 0, 7 and 14 post-immunization. Antigen-specific IgG are significantly reduced in TRPV1-Cre/DTA mice as compared to wild-type mice for 28 days (anti-KLH IgG U/mL, D14: wild type, 23,00,803 ± 7,24,085, n=4 versus TRPV1-Cre/DTA, 71,481± 35,903, n=7, *** p< 0.001 and anti-KLH IgG U/mL, D28: wild type, 17,96,135 ± 5,09,461, n=4 versus TRPV1-Cre/DTA, 88,633 ± 44,366, n=7, ** p< 0.01; **Figure 1A**). Next, we assessed antibody responses to hapten coupled to a carrier protein. We used OVA as the carrier protein and (4-hydroxy-3-nitrophenyl) acetyl (NP) as the hapten. Naive control and TRPV1-ablated mice wrere immunized with 12.5 μg of NP-OVA. Mice were bled at 7,14 and 28 days after the immunization. Anti-NP specific IgG antibody responses were measured by ELISA using NP_2_-BSA for antibody capture because high-affinity antibodies bind to proteins derivatized with small numbers of NP groups, whereas low-affinity antibodies do not (Reth et al., 1978). High titers of high-affinity anti-NP antibodies are found in control mice immunized with NP-OVA at day 28 (anti-NP_2_ IgG_1_ AU, D28: wild type, 18,401 ± 3,393, n=18). In contrast, serum titers of high affinity anti-NP antibodies (**Figure 1B**) are significantly decreased in TRPV1-ablated mice (TRPV1-Cre/DTA, 9,028± 1,498, n=18, ***p< 0.001). Together, these results indicate that TRPV1 nocioceptors are required for competent antigen-specific antibody responses.

**Figure 1.**
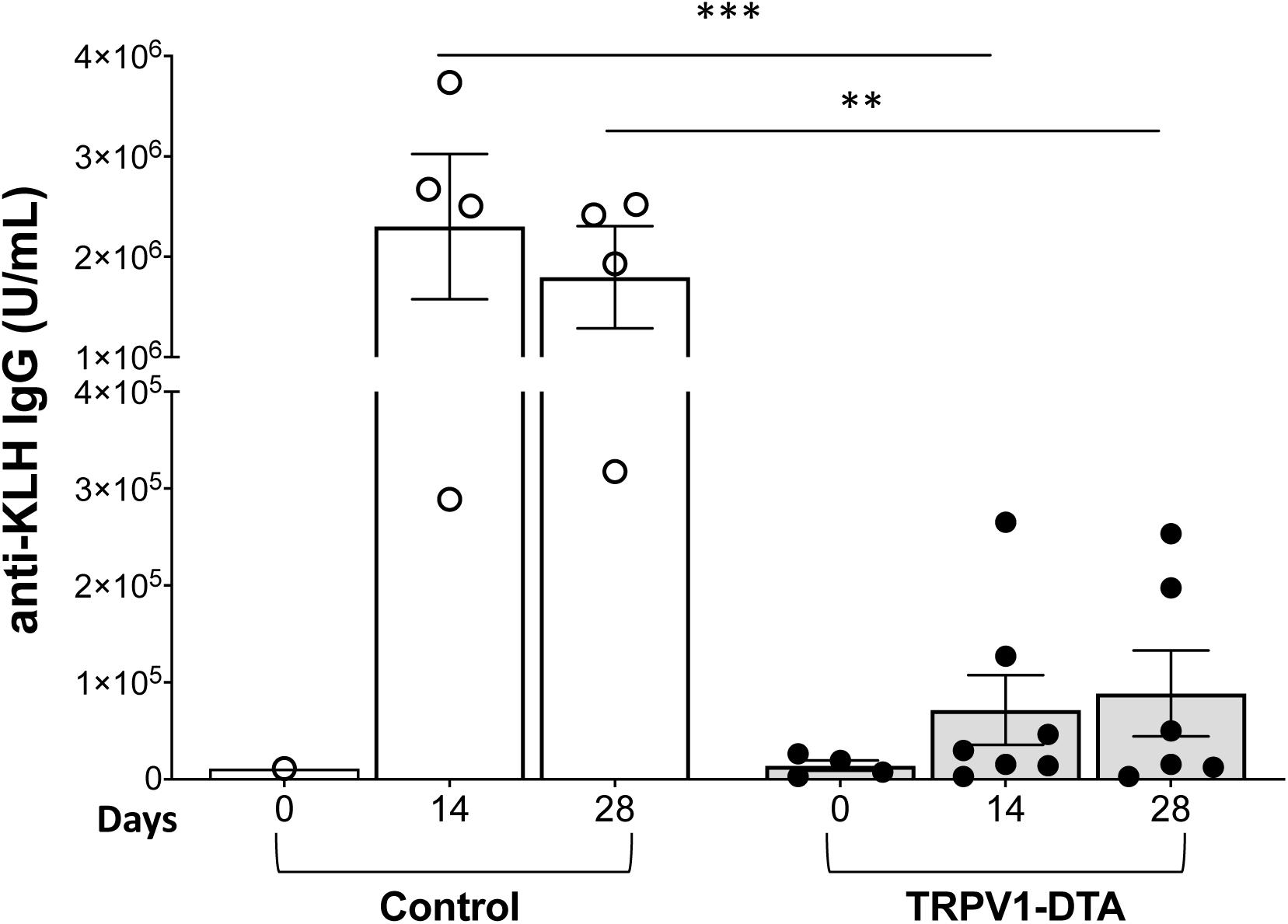
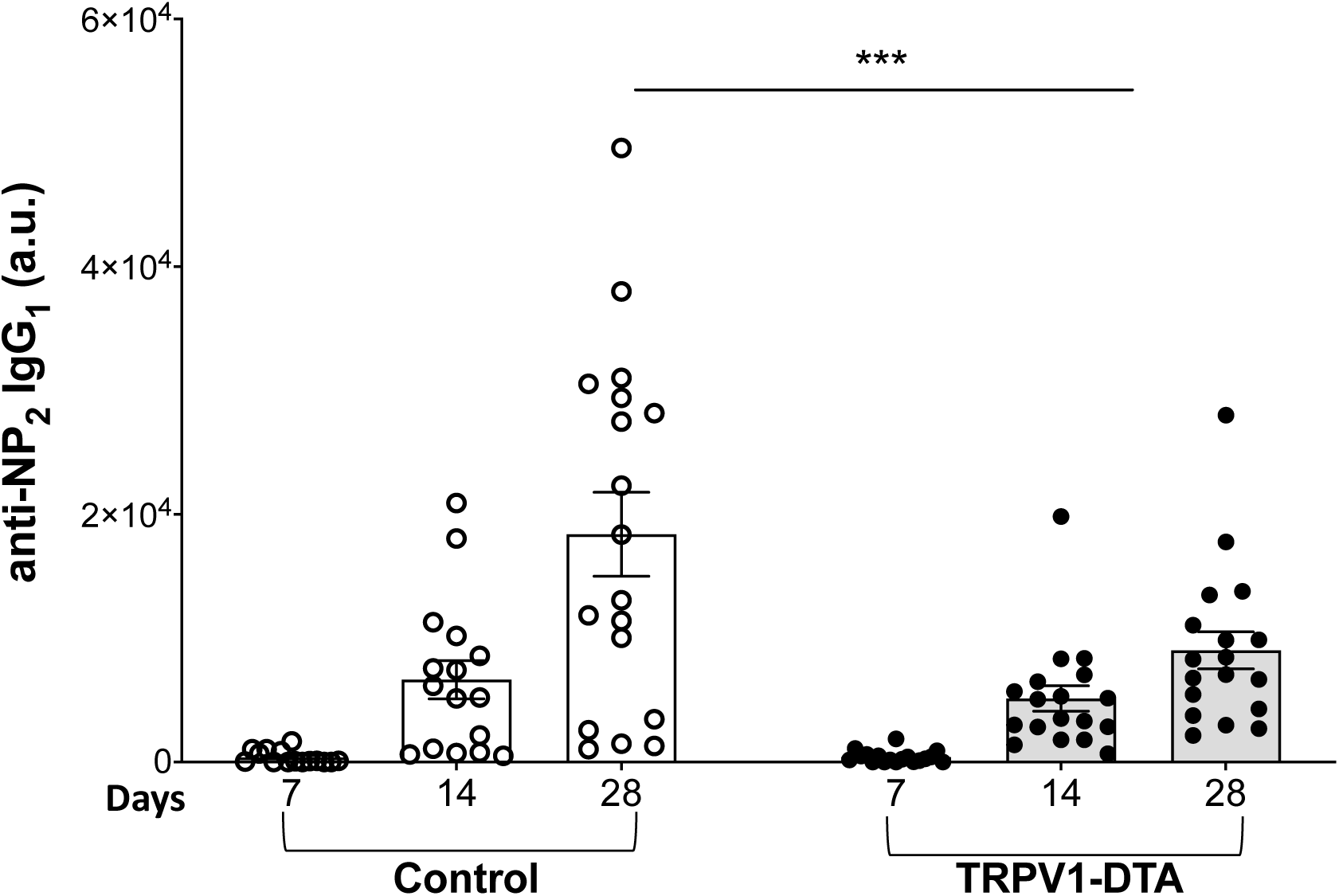
TRPV1 ablation impairs antigen-specific antibody response following immunization. **(A)** TRPV1-DTA (n=7) or wild type mice (n=5) were immunized with 4 mg/kg KLH intraperitoneally and serum collected every 7 days. Serum was assayed for anti-KLH IgG antibodies by ELISA. Data is represented as individual mouse data point with mean ± SEM. Ordinary one-way ANOVA with multiple comparisons test between groups: wild-type versus TRPV1-DTA; **p<0.01, ***p<0.001. TRPV1-DTA mice or wild-type mice were also immunized with NP-OVA in the foot pad and serum collected every 7 days. **(B)** High affinity NP-specific IgG1 antibodies were quantified by ELISA. Data is represented as individual mouse data point with mean ± SEM. (n=18/group from 2 repeated experiments). Two-way ANOVA with multiple comparisons test between groups: wild-type versus TRPV1-DTA; ***p<0.001.

### TRPV1 ablation does not reduce T and B lymphocyte numbers or impair B cell class switching

TRPV1 is expressed in T cells, macrophages, dendritic cells and mast cells (Helyes et al., 2007; Razavi et al., 2006; Szabó et al., 2005; Kimball et al., 2004; Collier, 2001), so it was possible that DTA expression in these mice may have reduced T and B cell numbers. To address this possibility, total TRCβ+ T cells and B220+ B cells were counted in the spleen of wild-type and TRPV1-ablated mice. There were no significant differences in the frequency of B cells (**Figure 2**) (wild type, 54.91% ± 3.389%, n=8 versus TRPV1-Cre/DTA, 54.89% ± 7.42%, n=10) or T cells (**Figure 2**) (wild type, 28.38% ± 9.772%, n=8 versus TRPV1-Cre/DTA, 29.2% ± 10.52%, n=10) in TRPV1-ablated mice. Thus, TRPV1-Cre/DTA mice have equivalent numbers of T and B cells required to generate a primary immunization response.

**Figure 2.**
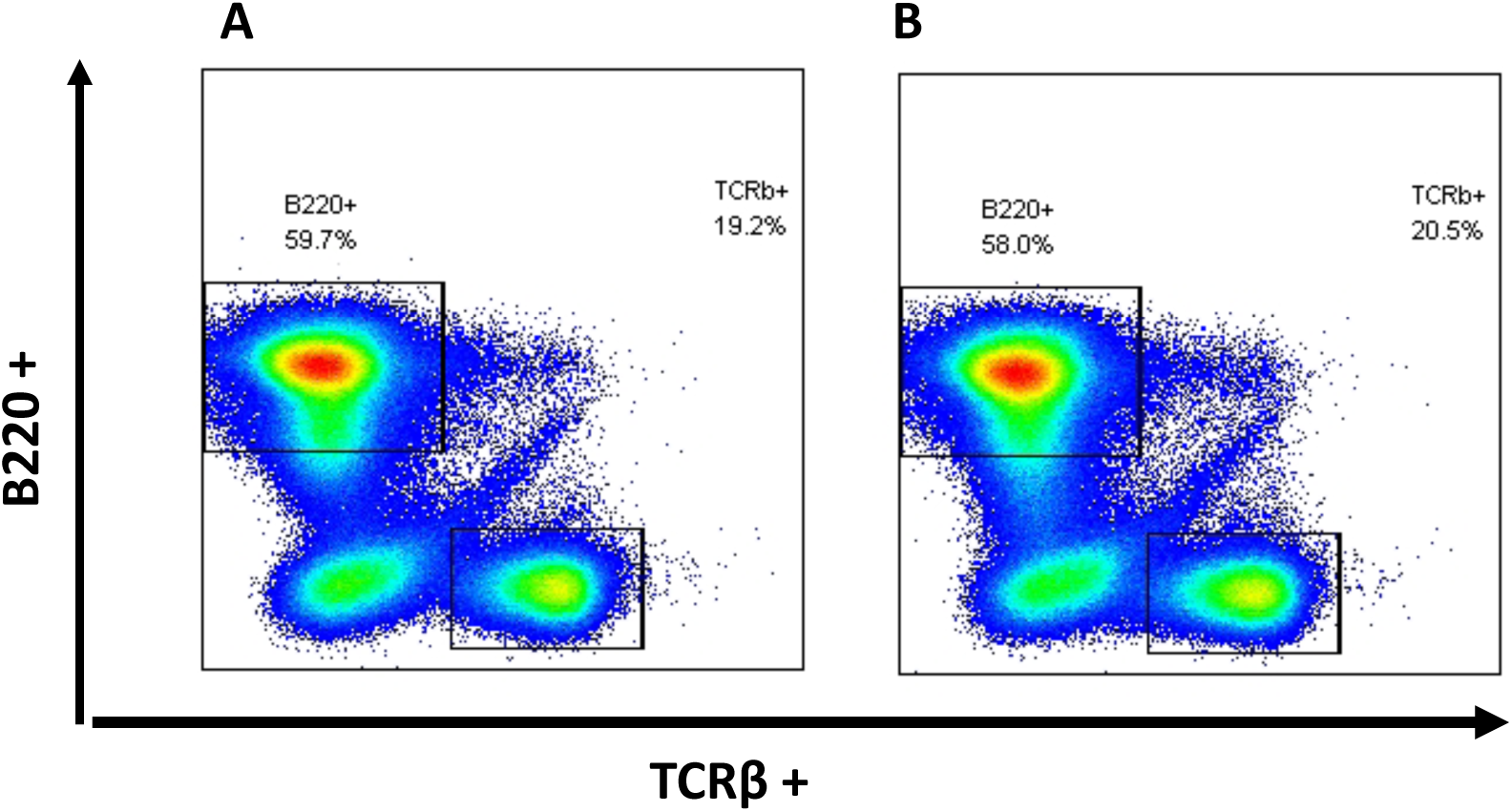
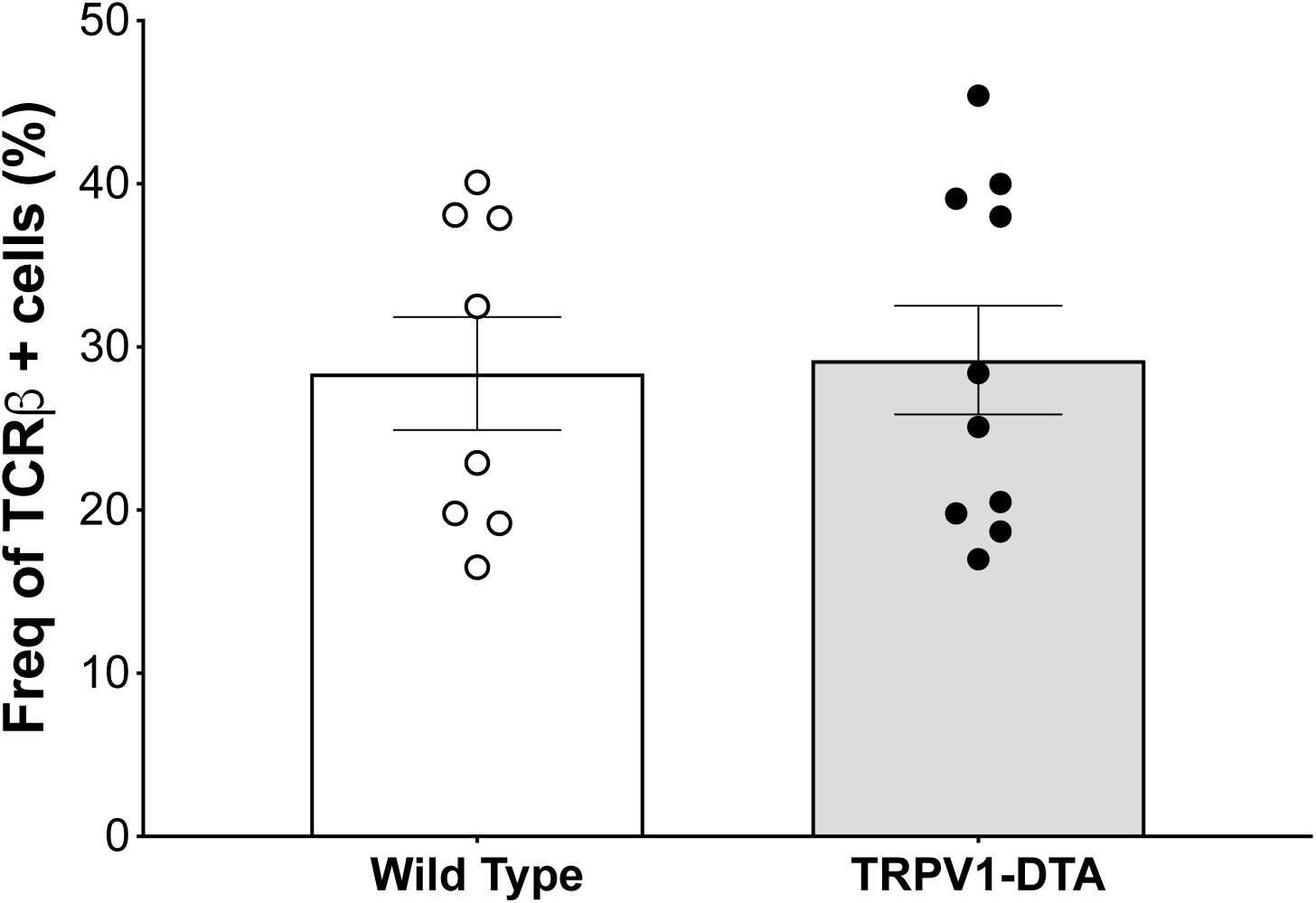
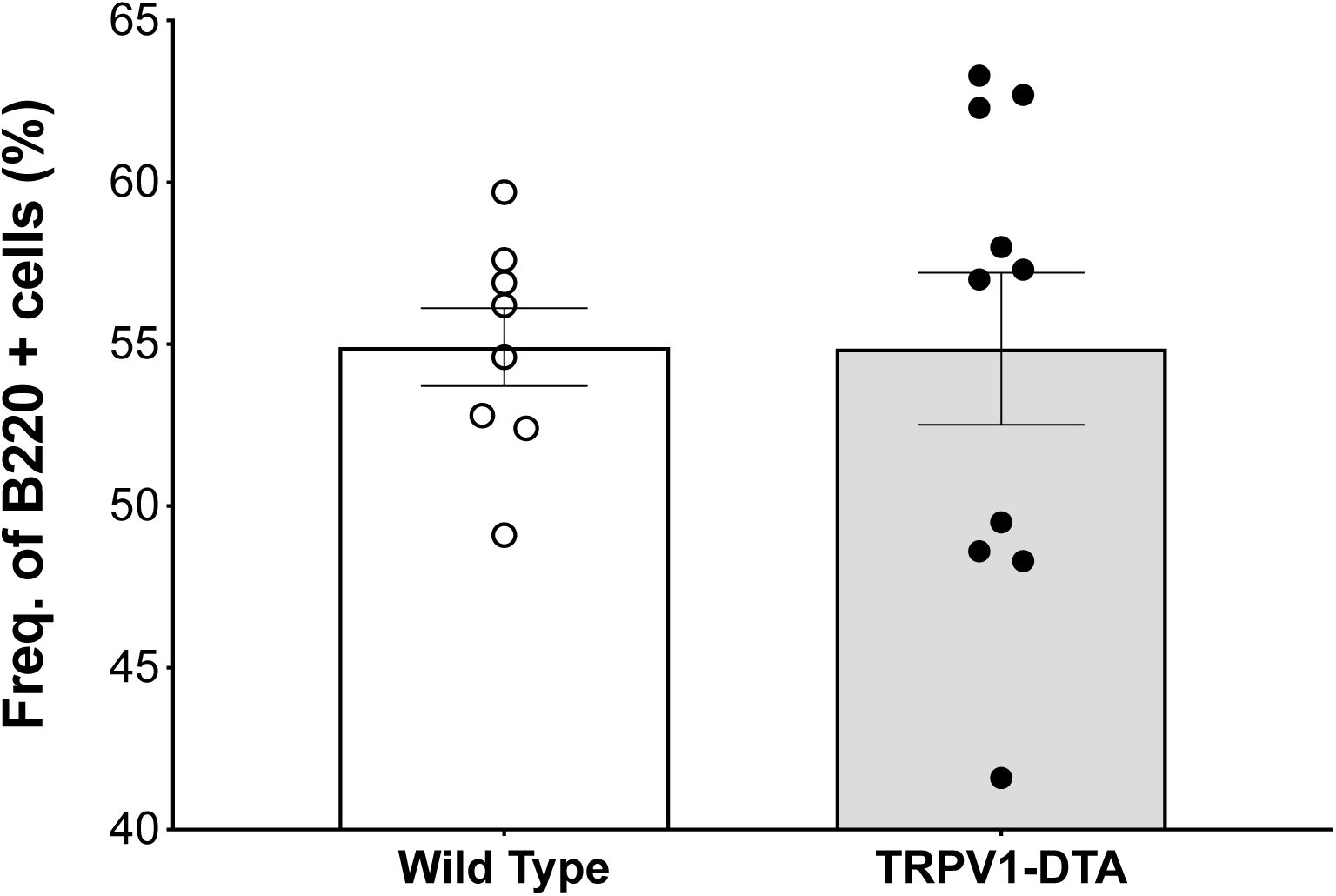
TRPV1 ablation does not change the frequency of T or B Lymphocytes in comparison to the wild-type mice. Total T cell and B cell numbers were quantified in spleens isolated from wild-type (n=8) and TRPV1-DTA (n=10) mice by flow cytometry. Representative scatterplots showing B220+ cells versus TCRβ + cells from **(A)** wild-type and **(B)** TRPV1-DTA mice. Data is representative of 4-5 animals per group. The frequency of **(C)** TCRβ + cells or **(D)** B220 + cells revealed no difference between wild-type and TRPV1-DTA mice. Data is represented as individual mouse data point with mean ± SEM. (n=8-10/group from 2 repeated experiments).

Having confirmed that TRPV1-ablated mice do not have impaired T and B cell numbers, we next asked whether B cells harvested from TRPV1-ablated mice are capable of producing a functional response when challenged. Thus, we cultured total splenocytes from naïve wild-type and TRPV1-ablated mice on ELISpot plates specific for murine IgG to assess the number of antibody secreting cells (ASC) (**Figure 3A**). The cells were challenged with lipopolysaccharide (LPS) to induce an antibody response. LPS, a component Gram-negative bacterial cell wall, activates B cells by way of innate toll-like receptors (TLRs) 4 and/or TLR2 signaling pathways (Akira and Takeda, 2004). Interestingly, no significant difference in the number of antibody secreting cells is observed in splenocytes from TRPV1-ablated mice as compared to wild-type mice (**Figure 3A-B**; IgG^+^ASC/2 x 10^5^ cells, Wild-type mice: 111 ± 19 versus TRPV1-DTA: 113 ± 14, n=3 experiments each with n=3 replicates each). Moreover, comparable levels of secreted IgG are observed in the cell supernatants of wild-type and TRPV1-ablated mice (**Figure 3C**) (ng/ml IgG, control, 7.8 ± 4.7 versus TRPV1-DTA, 7.8 ± 6.6, n=4 experiments each with n=3 replicates each).

**Figure 3.**
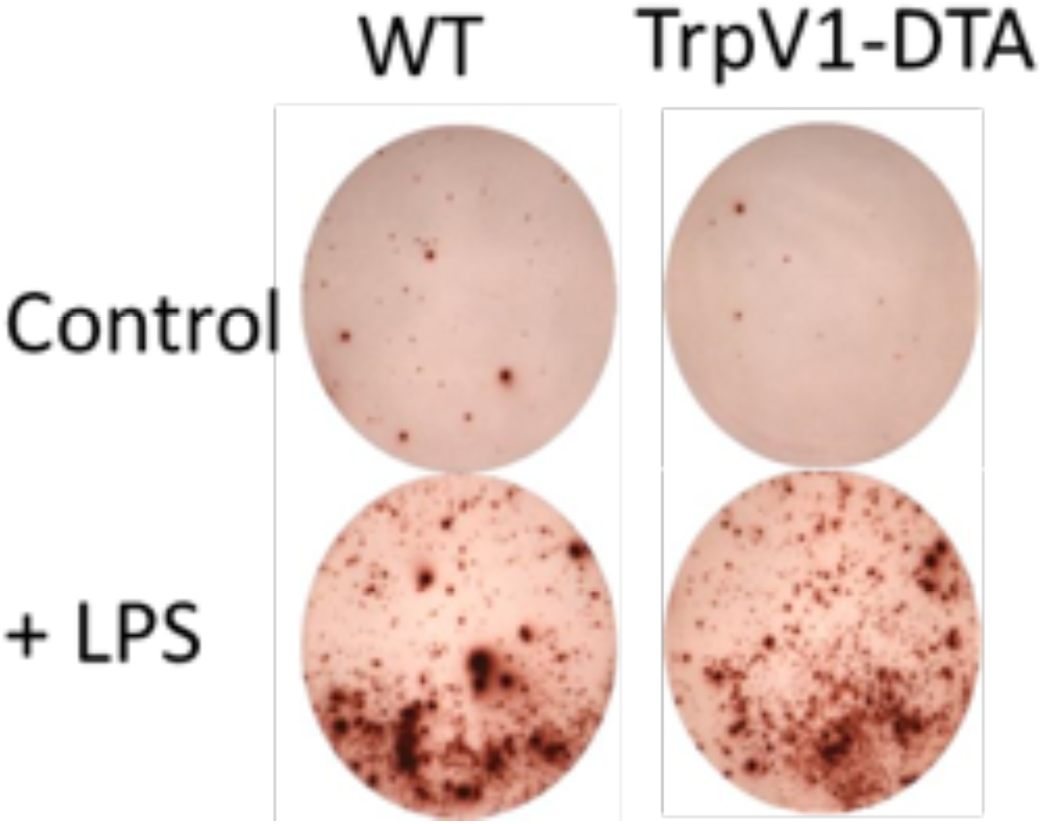
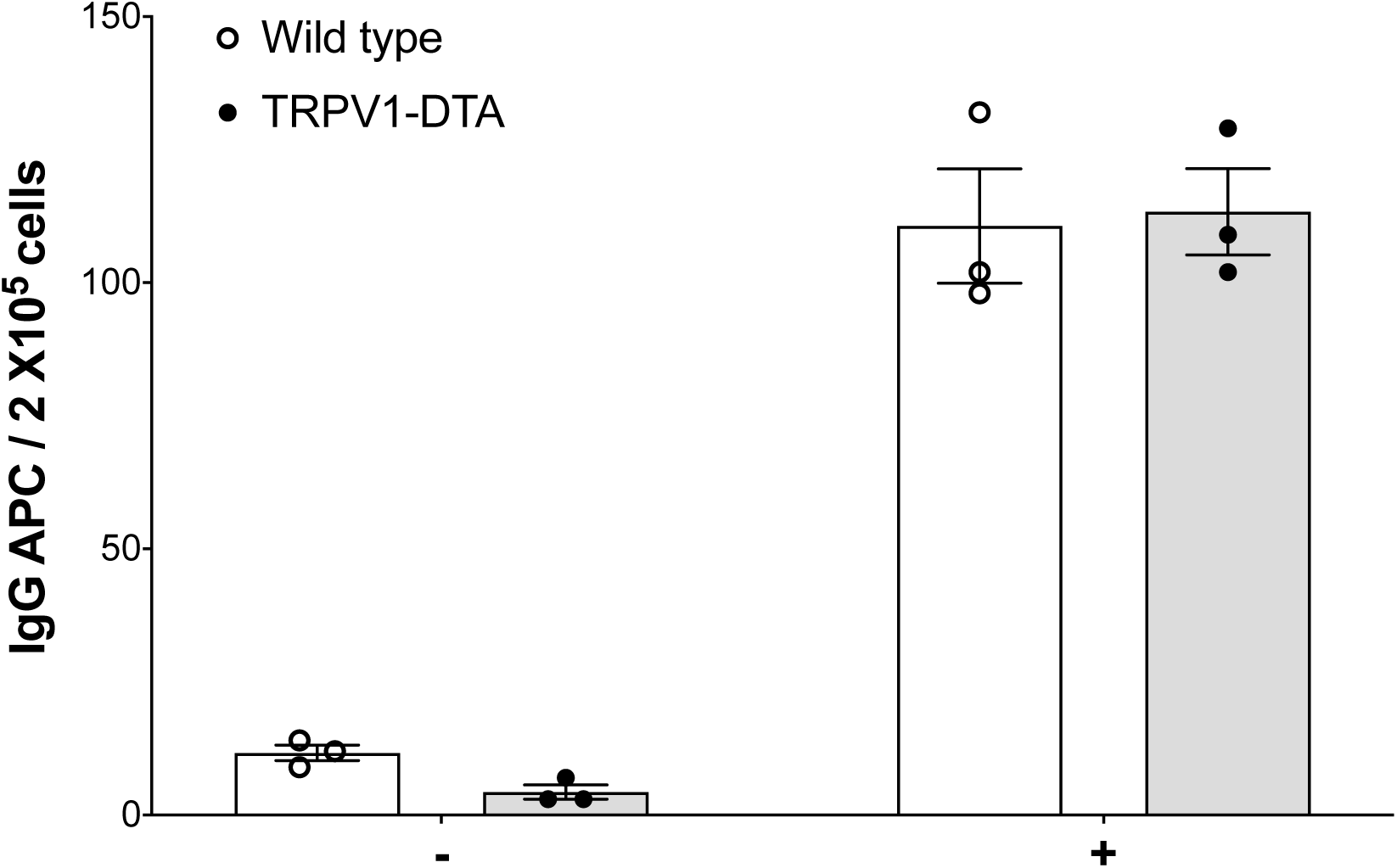
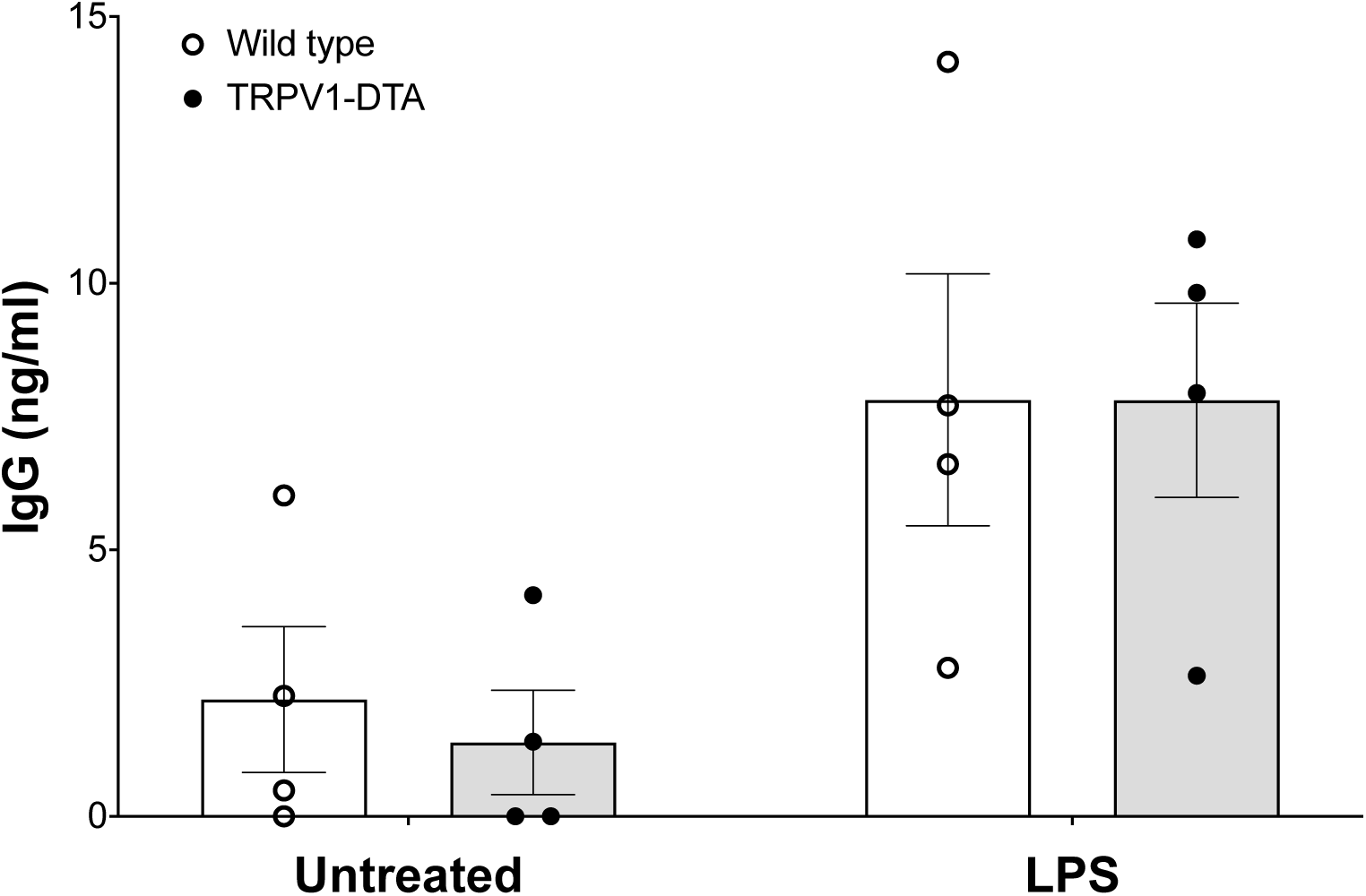
TRPV1 ablation does not impair the antibody response in *in vitro* activated splenocytes. Splenocytes from wild-type or TRPV1-DTA mice were stimulated *in vitro* with LPS, and the number of antibody secreting cells (ASC) was determined by ELISpot assay on anti-IgG coated plates. (**A**) Representative images from total IgG ELISpot assay performed on splenocytes from wild-type mice or TRPV1-DTA mice. Data is representative of 3 independent experiments. **(B)** Quantification of ASCs in unstimulated and LPS-stimulated splenocytes from wild-type and TRPV1-DTA mice using anti-IgG ELISpot assay. **(C)** Supernatant from unstimulated or LPS-stimulated splenocytes from wild-type and TRPV1-DTA mice was collected after 72 hours. Levels of total IgG secreted was measured using ELISA. Data represented as as individual data point for each experiment with mean ± SEM. Each experiment was carried out with at least 3 replicates per experiment.

### Optogenetic stimulation of TRPV1-expressing neurons enhances antigen specific antibody responses

In addition to sensory input for stretch and pain, TRPV1-expressing nociceptor neurons also exert local and systemic effector functions (Szabó et al., 2005). Next, to establish a causal role for TRPV1+ neurons in enhancing antigen-specific antibody responses, we used an optogenetic approach to selectively activate TRPV1-expressing neurons during immunization. This enabled us to assess the potential TRPV1-expressing nociceptor to immune cell signaling axis without *a priory* knowledge of the corresponding molecular or signaling mechanisms. The advances in optogenetics enables integration of optics and genetics to precisely track and control neuronal activity in living animals. Specifically, optical control of a neuron is achieved by targeted expression of a light gated cation channel, e.g. channelrhodopsin-2 (ChR2), in a selective population of neurons, thereby allowing temporally precise modulation of targeted neuronal activity (Deisseroth, 2011; Zhang et al., 2010; Boyden et al., 2005). To specifically activate TRPV1-expressing neurons during immunization, we used TRPV1-Cre/ChR2 mice in which TRPV1 lineage neurons expressed ChR2, generated by crossing TRPV1-Cre mice with Rosa26^ChR2-eYFP/+^ (ChR2) mice.

To directly test changes in antigen-specific antibody responses induced by ChR2-mediated TRPV1 excitation, we optically stimulated dorsal paw innervating TRPV1-expressing sensory neurons in TRPV1-Cre/ChR2 mice with blue light (450-490 nm, 3Hz, 20% DC, 4.7mW, 15 min) directed through a LED source during antigenic challenge (**Figure 4A**). C57BL/6 mice devoid of ChR2 were exposed to identical stimulation parameters to control for the light exposure procedure. Following light stimulatiion and immunization, serum anti-KLH antibody levels were monitored in mice at day 14 and 28 post-immunization. Optogenetic stimulation of TRPV1+ neurons significantly increased antigen-specific antibody responses in TRPV1-Cre/ChR2 mice as compared to wild-type mice (anti-KLH IgG ng/ml, TRPV1-Cre/ChR2: 2,581 ± 1,165, n= 6 versus Control, 1,224 ± 939, n=8; * p<0.05) (**Figure 4B**). Together, these findings describe for the first time that TRPV1+ nerves mediate the development of antigen-specific antibody responses in mice.

**Figure 4.**
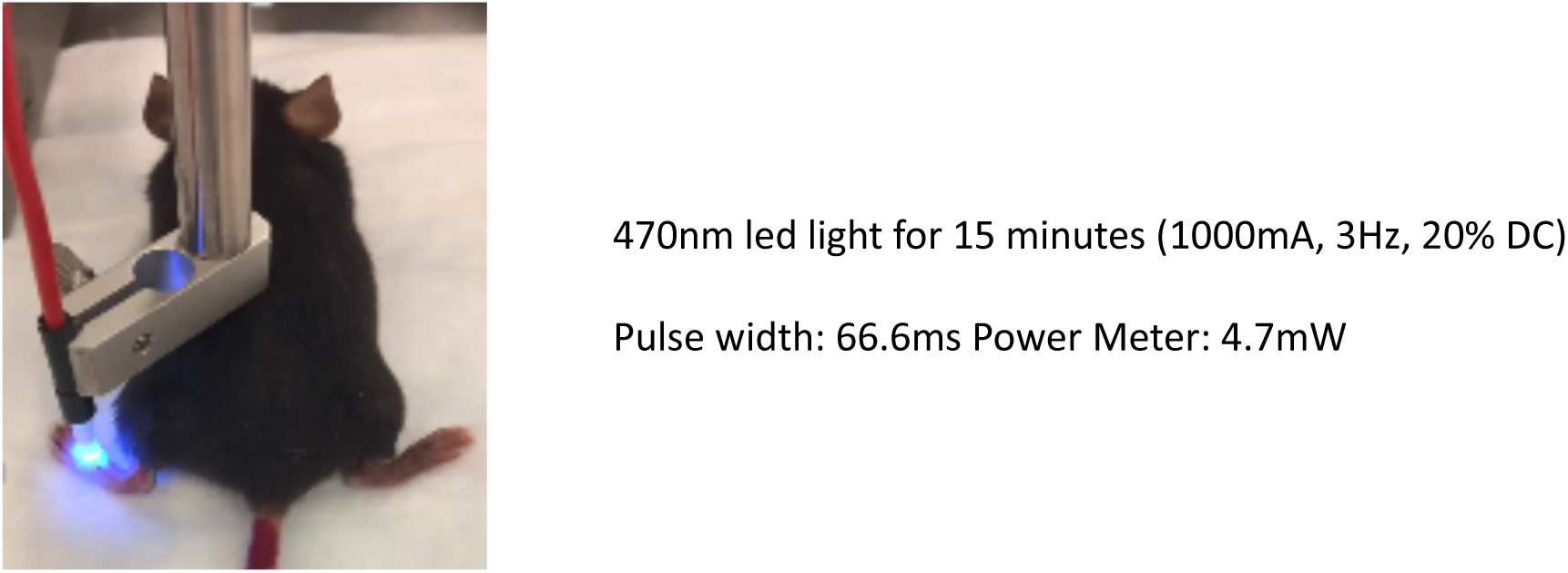
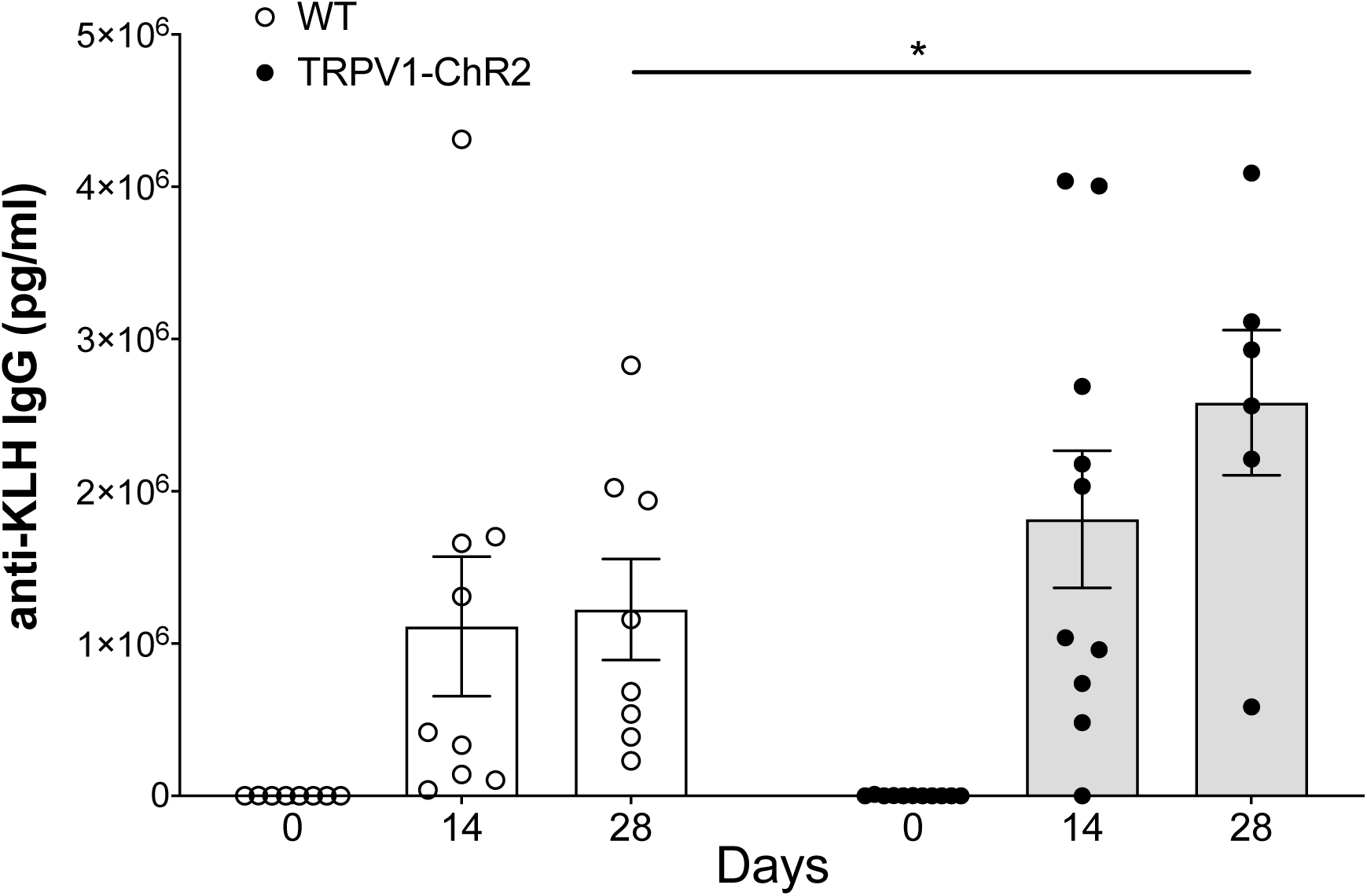
Selective optogenetic activation of TRPV1-expressing neurons enhances antigen-specific antibody response following immunization. Optogenetic stimulation of TRPV1-expressing neurons in the DMN attenuates serum TNF in endotoxemic mice. TRPV1-ChR2 mice (n=11) or wild-type mice (n=9) were subjected to optogenetic stimulation using blue light (473 nm, 3Hz, 20% duty cycle, 15 minutes, 1000mA) on the dorsum of the hind paw. **(A)** Representative image of the positioning of the 470 nm light source used to stimulate the dorsum of the hind paw. **(B)** Mice were immunized with 3 mg/kg KLH on the same paw immediately following optogenetic stimulation, and serum collected every 7 days. Serum was assayed for anti-KLH IgG antibodies by ELISA. Data is represented as individual mouse data point with mean ± SEM. Two-way ANOVA with multiple comparisons test between groups: wild-type versus TRPV1-ChR2; *p<0.05.

## Discussion

The nociceptor neurons and immune cells work in concert to protect the organism from environmental dangers. Although the interactions between the immune system and nervous system are known to contribute to systemic immunity, how peripheral nerves regulate antigen-specific antibody responses remains unclear (Rosas-Ballina et al., 2011; Chiu et al., 2012a). Here, using genetic and optical tools, we present the first observations showing that selective ablation of TRPV1-expressing cells inhibits antibody production, and selective optogenetic activation of TRPV1 neurons stimulates antibody production. Whereas ablation of sensory TRPV1 nociceptors significantly reduces antigen-specific antibody formation, direct activation of local nociceptors with light promotes antibody response. Although sensory input mediated by exposure to painful heat was previously shown to enhance memory development in the brain (Wimmer and Buchel, 2015), it was previously unknown whether neural signals influence memory development in antibody producing cells. The present findings indicate that nociceptors mediate information storage about novel antigens as specific antibody responses in adaptive immunity.

Numerous studies have revealed that nociceptors are involved in mediating adaptive immune responses. Genetic or pharmacological disruption of TRPV1 activity attenuated airway injury and asthma features in murine models of allergic airway inflammation (Rehman et al., 2013; Mabalirajan et al., 2013), while TRPV1 denervation attenuated dendritic cell numbers in response to antigenic challenge (Kradin et al., 1997). Studies on experimental models of atopic dermatitis and psoriasis have demonstred that TRPV1-positive sensory neurons can potentiate adaptive immune responses (Riol-Blanco et al., 2014; Wilson et al., 2013). Administration of capsaicin to rat neonates decreased antigen-specific antibody secreting cells as observed by direct and indirect plaque assay methods (Helme et al., 1987a; b). Another study found that capsaicin administration reduced IgA and IgG synthesis in cultured lymphoid cells from aerosol immunized animals (Nilsson et al., 1991). However, in addition to stimulating TRPV1 neurons and lymphocytes, capsaicin, when administered in higher doses, is cytotoxic. Thus, the prior findings may be the product of TRPV1-mediated signaling, or from depletion of TRPV1 expressing cells. Here we show that TRPV1-expressing nociceptors are crucial to the generation of antigen-specific antibody responses. Our results that selective optogenetic stimulation of the nociceptors expressing TRPV1 enhances antibody responses gives direct evidence for nocioceptor signaling in the development of antibody-dependent memory.

In TRPV1-DTA mice studied, we also obtained evidence that the nociceptors, but not lymphocytes were depleted. Although mostly expressed in sensory neurons, TRPV1 has been shown to be expressed in T lymphocytes, monocytes, macrophages and dendritic cells (Bertin et al., 2014; Omari et al., 2017). However, it is important to note that in majority of these studies TRPV1 messenger RNA (mRNA) but not protein expression was analyzed and cell lines but not primary human or murine lymphocytes were used (Schwarz et al., 2007). Although we cannot exclude the possibility that the TRPV1 lymphocytes are somehow impaired in the DTA animals, together the evidence that 1) optogenetic stimulation of neurons enhances antibody response; 2) the numbers of T and B cells was not decreased in TRPV1-Cre/DTA animals; and 3) the B cells from the TRPV1-Cre/DTA animals maintain the capacity to undergo class switching indicates a role for TRPV1-expressing neurons in the onset of adaptive immunity. Together, this indicates sensory neurons are a critical mechanism that culminates in producing antigen-specific antibodies.

There are several important implications from these results. First, understanding that the nervous and immune systems have interlocking mechanisms in memory formation extends a new theory for analyzing and studying immunity. The encounter with a novel antigen and storage of memory is not managed solely by the dendritic cells, macrophages and lymphocytes, but also by nociceptors. It is now interesting to consider how the sensory nervous system contributes to storing information in B cells about antigen encounters. Second, understanding that specific stimulation of sensory neurons by optogenetic technology enhances antibody responses offers tantalizing clues about how the quality of immunization may be enhanced by targeting nociceptors. Because adjuvant administration mediates painful inflammation and necrosis, it is now important to revisit the roles of nociceptor signaling in adjuvant mechanisms of immunity. Accordingly, we postulate an important role for the sensory nervous system in mechanisms controlling antigen specific antibody responses.

## Materials and Methods

### Animals

All procedures with experimental animals were approved by the Institutional Animal Care and Use Committee and the Institutional Biosafety Committee of the Feinstein Institute for Medical Research, Northwell Health, Manhasset, NY in accordance with the National Institutes of Health Guidelines. Animals were maintained at 25°C on a 12-hour light-dark cycle with free access to food and water. C57Black6/J, *R26*^LSL-DTA^ (B6.Cg-*Gt(ROSA)26Sor^tm2.1(CAG-EGFP,-DTA*G128D)Pjen^*/J), *R26*^LSL-ChR2^ (B6.Cg-*Gt(ROSA)26Sor^tm32(CAG-COP4*H134R/EYFP)Hze^*/J) and TRPV1-Cre+ (B6.129-*Trpv1^tm1(cre)Bbm^*/J) mice were purchased from Jackson Laboratory (Jackson Laboratory, Bar Harbor, ME, USA) and maintained as homozygous in fully accredited facilities at the Feinstein Institute for Medical Research. TRPV1-Cre mice were crossed to *R26*^LSL-DTA^ mice or to *R26*^LSL-ChR2^ to generate TRPV1-DTA or TRPV1-ChR2 mice respectively. 6-12-week-old age-matched female and male mice were used for experiments.

### Immunizations

For systemic immunization, mice were immunized with 4 mg/kg of Keyhole limpet hemocyanin (KLH) (Millipore Sigma, MA, USA) in 200μl of saline by intraperitoneal injection. For local immunizations, mice were immunized with either 12.5 μg of NP_25_-OVA (LGC Biosearch Technologies, USA) in 25ul PBS with Imject™ Alum (ThermoFisher Scientific, MA, USA) (2:1 ratio) in both hind paws or with 3 mg/kg KLH on one hind paw that was exposed to optogenetic stimulation. Blood samples were collected from immunized mice every 7 days, and levels of antigen-specific antibodies were determined by ELISA.

### Enzyme-Linked Immunosorbent Assay (ELISA)

Anti-KLH IgG were measured using commercially available ELISA as per the manufacturer’s protocol (Abnova, Taiwan). Total IgG was assayed using Invitrogen murine Total IgG kit (ThermoFisher Scientific). High-affinity NP-specific antibodies were measured by ELISA using 10 μg/ml of NP_2_-BSA as the coating reagent as previously described (Ersching et al., 2017). Briefly, serum was assayed in 4-fold dilutions starting at 1/100. NP-specific IgG1 was detected using a biotin conjugated rat anti–mouse IgG1 antibody (BD Biosciences, CA, USA) and developed with Strep-horseraddish peroxidase (R&D Systems, MN, USA) and tetramethylbenzidine (Sigma, MO, USA). OD450 was measured using a Tecan sunrise absorbance microplate reader (Tecan, Switzerland). Titers were calculated by logarithmic interpolation of the dilutions with readings immediately above and immediately below an OD450 of 0.3.

### Cell preparation

Spleens were isolated from wild-type or TRPV1-DTA mice and dissociated to a single cell suspension using a 70μm cell strainer. Red blood cells were lysed and the remaining cells used for culture experiments or flow cytometry.

### Flow cytometry

Total splenocytes were incubated in PBS containing 2% fetal bovine serum and 2 mM EDTA (Gibco, ThermoFisher, USA). Fc receptors were first blocked using unconjugated anti-CD16/32 prior to staining with the following antibodies: B220-FITC (RA3-6B2), TCRβ-BV510 (H57-597), BV650 (3/23). All antibodies were from Biolegend (CA, USA). Dead cells were excluded using LIVE/DEAD Fixable Near-IR Dead Cell Stain kit (ThermoFisher). All analyses were performed using FlowJo software (Treestar, USA).

### Cell Culture

Total splenocytes were cultured into 96 well tissue culture plates in triplicate at 2 x 10^5^ cells in RPMI 1640 (Gibco) supplemented with 1M NEAA, 1M Hepes, 1M Glutamax, 50μM βME2, 1% Pen/Strep and 10% FBS (all Gibco). Cells were stimulated with 500 ng/ml LPS (*Escherichia coli* 0111:B4, Sigma) for 72 hours. Supernatants were harvested for ELISA detection of IgG.

### Enzyme-Linked Immunosorbent Spot (ELISpot)

Total splenocytes were cultured on anti-IgG ELISpot plates and developed using the standardized ELISpot kit (Mabtech Inc., Cincinnati, OH, USA). Briefly, 96 well flat bottom multi-screen filter plates (Millipore, Billerica, MA, USA) were coated with 100 µL per well of anti-mouse IgG antibody (15 µg/ml). The plates were incubated overnight at 4°C, washed with PBS, and blocked using complete RPMI medium. Splenocytes were cultured at 2 x 10^5^ cells in triplicates in the plates overnight at 37°C with 5% CO_2_. ELISpots were developed as per the manufacturer’s protocol, scanned and analyzed at Cellular Technology Ltd. (Shaker Heights, OH, USA).

### Optogenetics

To activate TRPV1-expressing neurons by light pulse (473nm; power 4.7 mW; frequency 3Hz; 20% duty cycle, pulse width 67ms), a LED driver (Thor Labs, Newton, New Jersey) was connected to a blue LED source (Thor Labs). Parameters previously shown to demonstrate hypersensitivity in the paw for up to 24 hours were used (Daou et al., 2013). The power from the LED was measured using a power meter with the LED positioned at the same 1 cm distance from the sensor at the start and end of the experiment. The power measured from these settings at both points was 4.7mW and the pulse width was 67ms. The LED was positioned over the right hind paw at 1 cm distance from the skin (**Figure 4A**). TRPV1-DTA mice or C57Bl6 mice were stimulated with blue light on the right hind paw for 15 minutes. Immediately following optogenetic stimulation, mice were injected with 3 mg/kg of KLH subcutaneously in 20 μL into the same paw. Blood was collected weekly, and serum antibody titers tested.

### Statistics

Data were analyzed using Graphpad Prism 7 software using two-tailed unpaired Student’s *t*-tests or 2way ANOVA with Sidak’s multiple comparisons test or 1way ANOVA with Dunn’s multiple comparisons test. For all analyses, *P*≤0.05 was considered statistically significant.

## Author contributions

SSC and KJT conceived and designed the experiments; AT, TT, MG, and IS performed the experiments; AT, TT and SSC analyzed and interpreted data; AT, SSC and KJT wrote the manuscript; TWM, UA and PO contributed to finalizing the manuscript. All authors read and approved the final manuscript.

## Acknowledgments

The authors are grateful to James Rothman for discussion and advice throughout the study. This study was supported by grants from the National Institute of Health (NIGMS 1R01GM132672-01 to S.S.C., NIGMS 1R35GM118182-01 to K.J.T., and NIAID 1P01AI102852-01A1 to K.J.T. and S.S.C.).

## Availability of data and materials

The datasets used and analyzed during the current study are available from the corresponding authors upon reasonable request.

## Ethics approval

All experiments involving live animals were carried out in accordance with the National Institutes of Health guidelines for the use of experimental animals and were reviewed and approved by the Institutional Animal Care and Use Committee at the Feinstein Institute for Medical Research

## Abbreviations

TRPV1: Transient Receptor Potenial Vanilloid 1
KLH: Keyhole limpet hemocyanin
NP: 4-Hydroxy-3-nitrophenylacetyl hapten
fMRI: Functional Magnetic Resonance Imaging
CGRP: Calcitonin Gene-related peptide
ELISA: Enzyme-Linked Immunosorbent Assay
Ig: Immunoglobulin
DTA: Diphtheria toxin fragment A
ChR2: Channelrhodopsin-2
PBS: Phosphate Buffered Saline
OVA: Ovalbumin
BSA: Bovine Serum Albumin
EDTA: Ethylenediaminetetraacetic acid
NEAA: Non-essential Amino Acids
FBS: Fetal Bovine Serum
LPS: Lipopolysaccharide
ELISpot: Enzyme-linked Immunosorbent Spot
LED: Light-Emitting Diode
ANOVA: Analysis of Variance
TCRβ: T Cell Receptor Beta
ASC: Antibody Secreting Cell
Hz: Hertz
DC: Duty Cycle

